# A ‘capture-handling-restraint protocol’ elicits short-term stress responses in wild great tits (*Parus major*) but has little impact on reproductive success

**DOI:** 10.64898/2026.03.17.712382

**Authors:** Franziska Fröhlich, Lucia Mentesana, Caroline Deimel, Michaela Hau

## Abstract

Capturing and handling wild animals is essential for ecological and evolutionary research, yet their effects on physiology, behaviour, and reproductive success remain poorly understood. We investigated short-and longer-term consequences of a ‘capture-handling-restraint protocol’ in wild great tits (*Parus major*) over three breeding seasons. To assess short-term responses, we measured circulating corticosterone, a metabolic hormone that responds to unpredictable challenges, and automatically recorded provisioning behaviour. We also explored whether environmental and individual traits were related to provisioning latency (i.e., time to resume provisioning after capture). To evaluate longer-term effects, we monitored provisioning in the days following capture and related it to reproductive success (fledgling number and body condition). We predicted that longer handling would increase stress-induced corticosterone and provisioning latency, that these variables would be positively correlated, and that higher corticosterone and longer latencies would be associated with lower reproductive success. After capture, great tits showed elevated corticosterone and delayed provisioning. Contrary to our predictions, handling duration was negatively associated with stress-induced corticosterone in males (but not females) and did not affect provisioning latency. Provisioning latency was unrelated to corticosterone, environmental, or individual variables. Following capture, parents resumed provisioning, and short-term responses had little influence on reproductive success. We show that parental behaviour and physiology are affected by capture restraint protocols on the short term, but offspring condition and survival are not. However, these results should be interpreted cautiously, as our study lacks an uncaptured control group. Our findings highlight that evaluating welfare impacts requires rigorous study design incorporating both immediate and longer-term behavioural and fitness effects.

## 1. Introduction

As scientists, we bear responsibility not only towards research integrity but also towards the welfare of the animals we study. Ethical evaluations of research procedures are guided by the principles of the ‘Three Rs’ (Replacement, Refinement and Reduction; Russell & Burch, 1959), which were initially established for laboratory animals and later expanded into the ‘Nine Rs’ for studies involving wildlife (Curzer et al., 2013). The primary goal of these principles is to minimise harm experienced by animals used in research, as physiological and behavioural indicators suggest that animals perceive these procedures as negative affective states (Broom & Johnson, 2019; reviewed by Kaplan, 2022; Soulsbury et al., 2020). Negative welfare states can affect the validity, generalisation, and interpretation of research results (Poole, 1997; Prescott and Lidster, 2017). Moreover, when animals are subjected to research experiments during reproductive life-history stages, the effects of research activities might extend beyond the animal used for research and potentially also affect the well-being of their offspring.

Capturing, sampling, handling and restraining wild animals is a common practice in biological research (Boitani and Fuller, 2000; Duarte, 2013; Soulsbury et al., 2020). These human-animal interactions are conceptualised as eliciting negative experiences in the affected animals, resulting in physiological and behavioural changes that are mounted in response to this event that can be used as welfare indicators (Mellor et al., 2020). Yet our understanding of how these procedures affect the short-term behavioural responses of wild research animals remains incomplete (Gillies et al., 2020; Schlicht and Kempenaers, 2015; Soulsbury et al., 2020), particularly in relation to how these effects align with well-documented physiological responses. This is important because the application of the ‘Three Rs’, or rather ‘Nine Rs’ principles, relies on accurately documenting the adverse effects of procedures that animals involved in research experience.

The short-term physiological effects of ‘capture-handling-restraint protocols’ (Wingfield et al., 1982) are well documented in vertebrates. Upon encountering an unexpected environmental challenge (e.g., being captured by researchers), catecholamines are rapidly released from the adrenal cortex to mediate the Fight-or-Flight response (Romero and Butler, 2007). Even short periods of high adrenal activity can weaken the immune response, potentially increasing mortality (Broom and Johnson, 2019). As a second, slower response to a challenge, glucocorticoid levels begin to rise within three to five minutes (Wingfield et al., 1982). They typically peak between 30 and 60 minutes, reaching what is commonly referred to as ‘stress-induced glucocorticoid levels’ (reviewed by Romero, 2004; Romero & Reed, 2005). Since glucocorticoids can be more easily measured than the rapidly changing catecholamines, glucocorticoids are widely used as indicators of the physiological stress response (reviewed by De Kloet et al., 2008; reviewed by Dickens & Romero, 2013; Huber et al., 2021; Romero & Beattie, 2022; reviewed by Sapolsky et al., 2000; reviewed by Wingfield & Sapolsky, 2003). Elevated glucocorticoid levels (i.e., allostatic state) enhance the action of catecholamines and modulate physiological and behavioural responses, mediating allostasis, a process to maintain stability through change, helping the animal to cope with the stressor (McEwen and Wingfield, 2003). The cumulative cost of coping adds to the required energy (i.e., allostatic load). If the energy demand exceeds the energy supply, leading to an allostatic overload and resulting in an emergency life history stage, immediate survival can be promoted at the expense of functions such as reproduction (McEwen & Wingfield, 2003; Romero & Beattie, 2022; Romero & Wingfield, 2015; Wingfield et al., 1998; reviewed by Wingfield & Sapolsky, 2003). On the contrary, the short-term behavioural effects of ‘capture-handling-restraint protocols’ are poorly documented in vertebrates. For example, blue tits (*Cyanistes caeruleus*) exposed to this protocol during the breeding period stayed away much longer from the nest after being released (on average 4.2 hours) compared to their normal intervals between nest visits (Schlicht and Kempenaers, 2015). Although this study did not examine the relationship between short-term behavioural changes and physiological parameters, these behavioural responses seem to be contingent on physiological responses (i.e., the activation of the hypothalamic-pituitary-adrenal (HPA) axis and subsequent increases in glucocorticoids).

During the breeding season, short-term effects of ‘capture-handling-restraint protocols’ may also lead to longer-term consequences by altering parental behaviour, thereby indirectly affecting offspring development and survival. Experimentally induced short-term physiological stress responses have been associated with reduced reproductive success. In pied flycatchers (*Ficedula hypoleuca*), experimentally increased glucocorticoid levels to stress-induced levels during the nestling period led to reduced feeding and fewer surviving fledglings (Silverin, 1986). In black-legged kittiwakes (*Rissa tridactyla*), an experimentally induced short-term increase in corticosterone, a type of glucocorticoid, during the chick-rearing period led to decreased prolactin levels, which were associated with reduced nest attendance and increased latency to return to the nest, which in turn significantly reduced breeding success (Angelier et al., 2009). Similarly, in mammals, elevated cortisol, another type of glucocorticoid, reduced maternal care in postpartum marmosets (*Callithrix jacchus*), which carried their infants less than control females (Saltzman and Abbott, 2009). However, there seems to be little evidence that intrinsic stress-induced increases in glucocorticoids following ‘capture-handling-restraint protocols’ negatively affect breeding success. For example, eastern bluebird (*Sialia sialis*) females exhibited increased corticosterone levels following a ‘capture-handling-restraint protocol’, but these did not affect fledging success (Lynn et al., 2010). Likewise, Lane et al. (2025) showed that incubating female song sparrows (*Melospiza melodia*) exhibited elevated glucocorticoid levels and longer intervals between nest visits following capture. Still, glucocorticoid levels did not explain the latency to return to the nest after the capture event. On the other hand, parental short-term behavioural stress responses to ‘capture-handling-restraint protocols’ may also lead to longer-term consequences affecting offspring condition and survival. Nestlings may have to wait unusually long for parents to return after capture and during this time they are exposed to adverse conditions that cause negative affective states, such as hunger and cold. This short-term exposure to adverse conditions can potentially negatively affect proper development and survival on the long term. These consequences may persist if parents return to the nest after the capture event but provide reduced care compared to their behaviour before capture. However, empirical evidence from wild populations addressing these effects remains limited. One notable example is the study by Schlicht and Kempenaers (2015), which found that although blue tits stayed away from their nests for extended periods after release, parents resumed normal feeding rates upon returning and no negative association with breeding success was detected.

Since a ‘capture-handling-restraint protocol’ can affect multiple welfare indicators over different time scales, studies assessing their short-and longer-term physiological and behavioural relationships and their links to reproductive success are greatly needed. For birds, despite their common use as model species in field studies, such research remains limited.

In this study, we therefore investigated short-and longer-term relationships of a ‘capture-handling-restraint protocol’ with physiological and behavioural traits in a wild population of great tits (*Parus major;* Figure 1A). We conducted this study during the nestling provisioning period across three consecutive breeding seasons (2020-2022). To do so, we captured adult great tits on days seven to twelve after their eggs had hatched. We subjected them to a standardised ‘capture-handling-restraint protocol’, which included blood sampling and is commonly used in wild bird research (Figure 1B). After catching an individual, we took a first blood sample within three minutes to measure baseline corticosterone levels. Then, we marked and tagged the bird to allow for the unequivocal identification and restrained it in a cloth bag for 20-30 minutes. After this, we took the bird’s morphometric measures and a second blood sample to assess stress-induced corticosterone levels. Finally, we released the bird back into its nestbox. We automatically recorded feeding behaviour and nest abandonment by equipping each adult great tit with RFID leg transponders (Figure 1C) and using RFID antenna devices located at the nest-box entrance (125 kHz, NatureCounters, UK; Figure 1D). To assess reproductive success, we measured nestling condition on day 15 or 16 post-hatching and determined the final number of fledglings at fledging.

**Figure 1.**
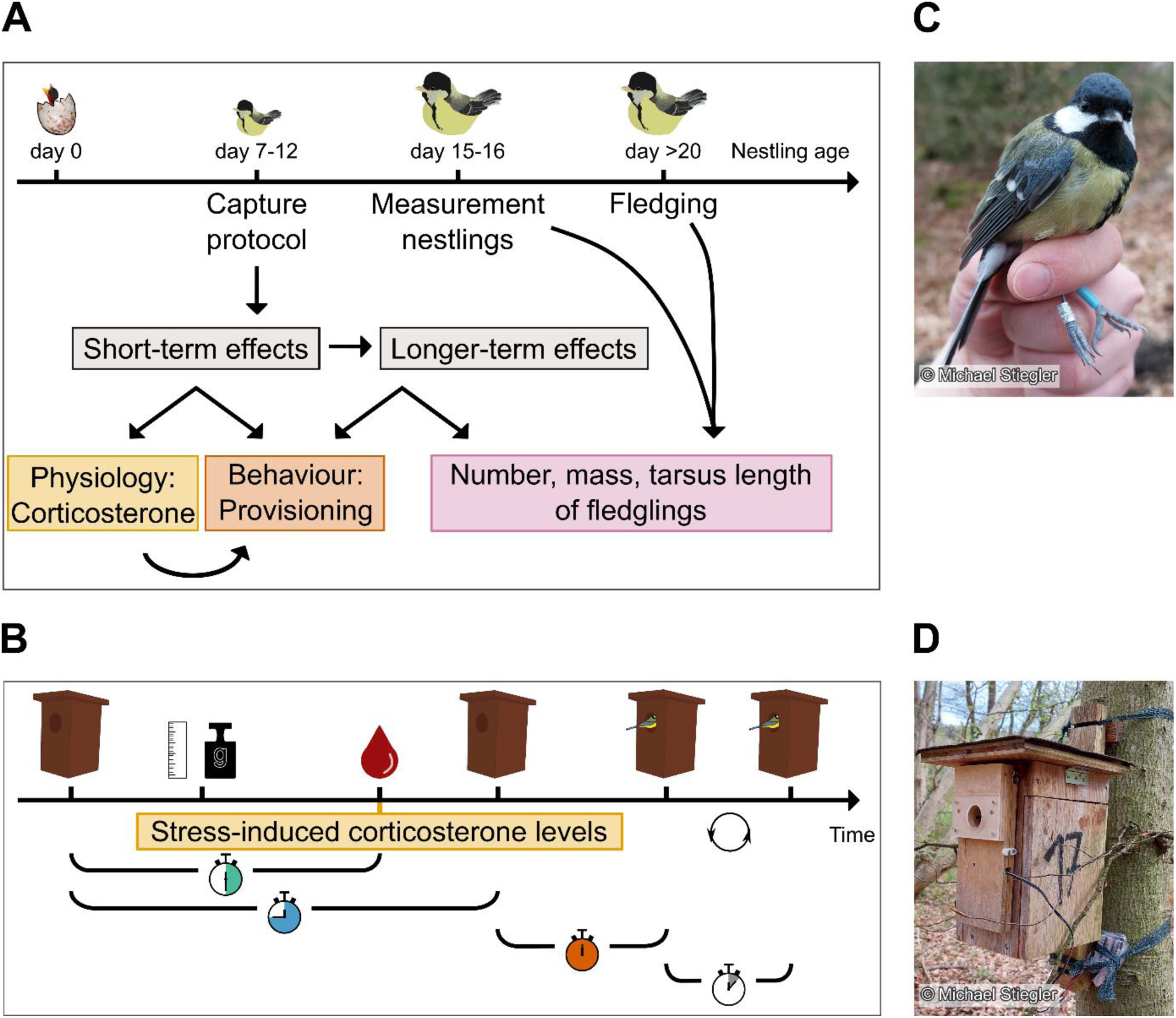
(**A**) Schematic representation of our research question that aimed at studying the combined short-and longer-term relationship of a ‘capture-handling-restraint protocol’ for breeding great tits. (**B**) Upon capture, we measured birds and extracted a blood sample to quantify the levels of baseline (not shown in the figure) and stress-induced corticosterone levels (green clock). Upon release (blue clock), we recorded provisioning latency (orange clock) and following nest visits (grey clock) using RFID transponders. (**C**) Great tit ringed with a numbered metal ring and an RFID transponder (blue ring). (**D**) Nest box with an RFID logger at the entrance hole.

First, we predicted that increases in handling duration would be correlated with higher stress-induced corticosterone levels, a greater likelihood of nest abandonment and increased time to resume provisioning (i.e., provisioning latency) after the catching event (Canoine et al., 2002; Monaghan & Spencer, 2014; reviewed by Romero, 2004). Because stress-induced corticosterone levels are suggested to support behavioural responses to cope with a stressor (reviewed by Andrews, 2022; Veen & Sivars, 2000; Wingfield et al., 1998), we predicted provisioning latency to increase with higher stress-induced corticosterone levels.

Secondly, an individual’s capacity to respond to the stress of capture and handling is contingent on its own body condition, health, life history traits and perceived trade-offs, as well as environmental mediators. Therefore, we examined whether provisioning latency was explained by ambient temperature at capture, parental condition or brood quality. For parental condition, we considered variables such as age, parasite infestation and mass and tarsus length, as these factors influence an individual’s allostatic load and therefore its capacity to sustain physiological and behavioural resources that are required for a response (McEwen & Wingfield, 2003; reviewed by Wu et al., 2025).

Thirdly, we examined if the ‘capture-handling-restraint protocol’ had longer-term relationships with behaviour (i.e., average time between two consecutive nest visits by a parent) and offspring condition (i.e., nest success, fledgling number, fledgling mass and tarsus length) and survival. Following (Schlicht and Kempenaers, 2015), we predicted that birds that returned to their nests after being captured would resume normal provisioning activity on the days following the capture. We also predicted that higher stress-induced corticosterone levels and longer provisioning latencies would be related to poorer nestling condition and decreased survival (Keller and Noordwijk, 1993; Monaghan and Spencer, 2014).

Lastly, because both male and female great tits care for the nestlings and nest success depends on contributions from both parents (Hinde, 2006; Norris, 1990), we examined whether nest abandonment by one parent would reduce fledging success. We predicted that the abandonment by one parent leads to a reduced fledging success, but not necessarily to complete nest failure.

## 2. Materials and Methods

### 2.1 Study site

We collected data for both first and second clutches during three consecutive breeding seasons (May-July of 2020-2022) in a nest-box population of wild great tits, located in the Ettenhofer Holz (48°03’26.7“N 11°15’11.0”E) in Upper Bavaria, Germany. Our study area contained 124 nest boxes and is a mosaic deciduous-coniferous forest of predominantly European beech (*Fagus sylvatica*) and Norway spruce (*Picea abies*), typical of managed forests in this area. This population has been the subject of a long-term study since 2013 and many individuals are fitted with a uniquely numbered metal leg ring (Hau et al., 2022).

### 2.2 Field Procedures

At the beginning of the breeding season, we visited the nest boxes every second day to determine when the first egg was laid, the final clutch size and when the first chick hatched (hatching date = nestling day 0, for more details regarding the study site and the standard nest box monitoring protocol (see Hau et al., 2022; Mentesana et al., 2024). When nestlings were between seven and twelve days old (mean ± SD = 8.56 ± 0.88 days), we caught the parents in the nest box between 07:00 and 14:00 hours. We took a blood sample (50-75 µl) from the brachial vein using a cannula needle (gauge size 26) and heparinised microhematocrit capillary for determining baseline corticosterone levels within three minutes after capture (mean ± SD = 118 ± 34 sec). We recorded the ID of the individual from the metal ring and RFID transponders, if present. Unmarked birds were fitted with metal rings and with a silicone leg ring containing a uniquely identifiable PIT tag (PIT Bird tag, EM4102 based EccelTechnology). Phenotypic sex was determined based on plumage characteristics and age (1^st^ year breeding vs. older) was estimated based on the colouration of the primary coverts. The age of birds that were ringed as nestlings was based on the year of birth. We proceeded with the ‘capture-handling-restraint protocol’ by placing the bird inside an opaque cloth bag for approximately 20 minutes (Wingfield et al., 1992, 1982). Then, the bird was taken out of the bag, morphometric measurements were taken (body mass to the nearest 0.25 g, tarsus and wing length to the nearest 0.1 mm) and the bird was checked for the presence of parasites (i.e., ticks, louse flies, feather mites). Next, we took a second blood sample (max. 25-30 µl) to determine stress-induced corticosterone levels and recorded the sampling time (i.e., the time between capture and finishing the second blood sample; mean ± SD = 28.70 ± 3.74 min, min. = 21.00 min, max. = 40.67 min). After the bleeding had stopped, we released the bird into its nest box through the entrance hole and again recorded the time to determine the total handling time (i.e., the time between capture and release; mean ± SD = 32.19 ± 5.07 min, min. = 8.00 min, max. = 45.00 min). Both blood samples were kept on ice until centrifugation in our laboratory. The plasma fraction was stored at-20°C until analysis (as described in Hau et al., 2022). Ambient temperature at capture was recorded automatically to the nearest 0.01°C every 30 min by a HOBO weather station (equipped with S-THB-M002 12 Bit Temperature sensors) located within the study area. We used the temperature recordings that were closest to our time of capture.

As proxies for nestling condition, we measured fledgling mass and tarsus length of nestlings shortly before fledging (97.7% on nestling day 15 and 2.3% on day 16). We checked nests daily for fledging and recorded if any nestlings had died in the nest since day 15 to determine the number of fledglings.

### 2.3 Corticosterone analysis

We quantified corticosterone levels as described in detail in Hau et al. (2022) using the Arbor Assays DetectX® Corticosterone Enzyme Immunoassay Kit with the high sensitivity protocol (Cat. No. K014). Before quantification, we conducted a double liquid-liquid extraction with diethyl ether of the plasma samples (Baugh et al., 2012; Hau et al., 2022; Ouyang et al., 2011). After evaporation of the diethyl ether, samples were reconstituted over night at 4°C in the assay buffer provided with the kit and assayed in duplicate on the following day. We included several control samples in each assay batch, which were taken through the entire extraction and analysis procedure. Two positive controls were included, containing stripped chicken plasma to which a known amount of a corticosterone stock solution was added, as well as two negative controls containing only ultrapure water. In total, we extracted and analysed 26 batches and plates (for further details on assay procedures, as well as intra-and inter-assay variation, see Supplementary Information 1). We measured two quality control (QC) samples on each plate, one at around 60% (QC 1) and one at around 30% (QC 2) percent binding. We measured both QC samples twice in duplicate on each plate and used these values to calculate intra-and inter-assay variation to evaluate extraction and assay consistency. Intra-assay variation was calculated by averaging the coefficient of variation between duplicate measurements of the control samples across all plates. Intra-assay variation was 13 ± 10% for QC 1, and 8.1 ± 4.5% for QC 2. Inter-assay variation was calculated by calculating the coefficient of variation across the corticosterone concentrations of the two control samples across plates. Inter-assay variation was 15.9% for QC1, and 9.1% for QC. Note that these values include not just variation on assay performance across plates, but also variation in extraction efficiency.

For nine individuals (three females, nine males), we measured unusually high corticosterone levels (range: 134.50-313.62 ng/ml) as compared to our long-term data from this population (Hau et al., 2022). Since these samples were processed together on the same plate, the values are likely not true biological measurements but rather the result of human error and were excluded from the analyses.

### 2.4 Automatic nest visit recordings

We used an RFID system (Radio Frequency IDentification) to automatically record the date and time of each bird’s visit to a nest box. Each nest box was equipped with a reading antenna, positioned around the entrance hole.

The reliability and consistency of RFID devices can vary significantly between studies (Hughes et al., 2021; Iserbyt et al., 2018; Schlicht and Kempenaers, 2015), making it essential to perform validations for each study. Therefore, we performed validation tests to assess the reliability of our setup. In 2020, we validated the RFID system by comparing its nest visit data with video recordings prior to the capture (for details see Supplementary Information 2). This allowed us to determine how accurately entries and exits were recorded and to estimate the time spent inside and outside the nest box, which is important for identifying separate nest visits. Entry registration was reliable, with 94.3% of entries correctly registered. Exit registration was much lower, with only 22.4% of exits properly registered because birds exited the nest boxes too quickly for the system to respond and record. Ninety per cent of the nest visits lasted less than one minute and 52 seconds. Based on these results, we excluded any consecutive RFID recordings by the same bird that occurred within a two-minute interval, as they likely belonged to the same visit (see Supplementary Information 2).

We conducted a second video validation in 2022 to test the performance of the RFID system following the release of birds into the nest box after capture. When we placed birds at the entrance hole, the entry was properly recorded in only 53.6% of cases. The registration of exits was 42.9%, likely due to birds exiting more slowly and cautiously. Birds also stayed much longer inside the nest box after the release, up to 40 minutes. Therefore, to calculate provisioning latency (i.e., the time between release and the bird’s first return to resume provisioning) and to avoid confusing the exit after release with the subsequent re-entry, we filtered the data to include only the first registered movement that occurred at least 51 minutes after release. This cutoff was based on the latest exit (after 40 minutes) and the earliest return (after 62 minutes) observed in the video analysis in 2022. Thus, 51 minutes was a conservative midpoint between these values. To filter the data, we created a timestamp variable using the ‘as.POSIXct’ function (R Core Team, 2024). Birds that abandoned their nests were excluded from the final analyses. For the birds that returned on the next day, the duration of the night was included in the calculation of provisioning latency.

One common error with these automatic recording systems is the registration of non-parental individuals, either due to incorrect PIT tag readings or because birds other than the social parents inspected the nest box. Non-parental IDs (e.g., IDs of other tagged birds, misread IDs) were recorded in ∼60% of the nests and were excluded from the analyses. Other errors were time-shifted RFID recordings (in ∼5% of the nests), due to failing batteries and automatic resetting of the logger clock. To detect such instances, we visually inspected the data for entries occurring outside the birds’ usual activity times (normal activity time: ∼04:30 to 21:30h), compared them with notes on battery changes and discrepancies between the logger clock and field-recorded notes and adjusted the data accordingly. If a time shift could not be resolved due to missing information on the correct time and it occurred around the time of the capture of a bird, the data were excluded from the analyses (this occurred for only one nest).

Since this study did not include a control group of uncaptured birds, we calculated the average provisioning interval (i.e., regular intervals between the nest visits during the two days before the capture) to better interpret provisioning latency. For this, we used data from birds that had already been equipped with a PIT tag during a previous capture (N = 83, females: 43, males: 40).

### 2.5 Statistical Analyses General Information

All statistical analyses were performed with the statistical software R (version 4.4.0, R Core Team, 2024) using the ‘*lm*’, ‘*lmer*’ or the ‘*glmer*’ function of the R-package ‘lme4’ (Bates et al., 2015). To ensure that model assumptions were met, we conducted a series of diagnostic checks by visually inspecting the residuals. All covariates were mean-centred to allow for comparisons of effect sizes across variables. We fitted statistical models using a Bayesian approach using the ‘*sim*’ function of the R-package ‘*arm*’ (Gelman and Su, 2021), to obtain 10,000 simulations for the posterior modes, from which we estimated the mean and the 95% Bayesian credible intervals (CrI). Following recommendations by Amrhein et al. (2019), we present our results by first describing their direction and then their statistical significance (defined as variables for which the 95% CI did not overlap with zero). To plot the results, we used the ‘*ggplot*’ function of the library ‘*ggplot2*’ (Wickham, 2016).

#### 2.5.1 Short-term relationship of the ‘capture-handling-restraint protocol’ with physiology and behaviour

To test for a relationship of our ‘capture-handling-restraint protocol’ with either physiology or behaviour, we ran three univariate linear mixed-effect models with a Gaussian distribution. Bird ID and nest ID were included as random factors. To test for the influence of our handling protocol on the birds’ physiology, we fitted stress-induced corticosterone levels as the response variable. Sampling time, date and time of capture were included as covariates. Sex (levels: females vs. males) was added as a fixed factor, as well as its interaction with sampling time. As random factors, we additionally included year and assay number. To test whether corticosterone levels increased in response to the protocol, we compared baseline (< 3 min) to stress-induced corticosterone levels using a two-sided paired t-test.

To test for a relationship between handling and the birds’ behaviour, we used log-transformed provisioning latency as the response variable. Handling time, date and time of capture were added as covariates. Sex was added as a fixed effect. The interaction between handling time and sex was also included. Additionally, bird ID, nest ID and year were included as random factors, but since year explained no variance, we removed it from the final model. For the interpretation of provisioning latency, we first compared the average provisioning interval one and two days before the capture. We performed two-sided paired t-tests to compare the average provisioning intervals between the two days preceding capture, analysing females and males separately to account for potential sex differences.

To test if our handling protocol led birds to abandon their nests (i.e., whether birds did not return to the nest after being captured), we ran a univariate generalised linear mixed-effect model with a binomial distribution fitting the return of a bird (levels: no vs. yes) as the response variable, handling time as a covariate, and sex as a fixed effect. We set the covariate handling time in interaction with sex. Nest ID and year were included as random factors.

To test for the influence of stress-induced corticosterone levels on the birds’ behaviour, we fitted the log-transformed provisioning latency as the response variable. Stress-induced corticosterone levels, date and time of capture were added as covariates. Sex was included as a fixed factor together with its interaction with stress-induced corticosterone levels. Additionally, we included bird ID and nest ID as random factors. We also included year as a random factor, but since it explained no variance, we removed it from the final model.

#### 2.5.2 Relationship of ambient temperature, parental condition and brood quality with behaviour

To test for the potential influence of other factors besides handling on provisioning latency, we performed five additional models with a Gaussian distribution and log-transformed provisioning latency as the response variable. We first investigated the influence of ambient temperature at capture by fitting this variable, date and time of capture as covariates, sex as a fixed factor and the interaction between temperature and sex. Bird ID and nest ID were included as random factors. Elevated temperatures may facilitate quicker energy replenishment for parents following a predation event due to increased foraging profitability (Avery and Krebs, 1984). According to this, higher temperatures could be expected to be associated with a lower provisioning latency.

Second, we examined whether the condition of the parents at capture influenced the time it took them to return to their nest box. For this, we ran two separate models for females and males, respectively. We included mass and tarsus length as covariates; age (levels: one year vs. older) and parasite infection (levels: no vs. yes) as fixed factors. For females, we included bird ID and year as random factors. However, for males, these variables did not explain any variance; thus, we ran a linear model. Parents in better condition might cope with stress more effectively than those in poorer condition. Conversely, individuals in poor condition might resume provisioning sooner, as their reduced prospects for future reproduction could favour greater investment in the current brood. Parental age might also influence provisioning latency. Older birds might have greater stress tolerance and respond less sensitively to capture, resulting in shorter provisioning delays (Monaghan and Spencer, 2014). Limited future mating opportunities might also drive them to suppress the behavioural short-term stress response of avoiding the nest after being captured at the nest and to return sooner to feed their offspring (Kitaysky et al., 1999).

Third, we tested for the association that brood quality (i.e., clutch size, offspring age and offspring body condition) might have with the parents’ behaviour. We ran two separate models for females and males. In each model, we added offspring age (median = 8 days, range: 7-12 days) and number of chicks (median = 6, range: 2-8) as continuous variables. We added clutch number (levels: 1 vs. 2) as a fixed factor. In both models, we included bird ID as a random factor. Solely for females, we also included year (because for males, it did not explain any variance). Brood quality may encourage parents to return to their nests more quickly, as offspring that are more numerous, older and in better body condition are likely to beg more loudly for food than offspring that are fewer, smaller and weaker (Bowers et al., 2019).

#### 2.5.4 Longer-term relationship of the ‘capture-handling-restraint protocol’ with proxies for reproductive success

To examine whether the protocol led to prolonged behavioural changes following capture, we analysed average provisioning intervals in birds that returned to their nests, separately for females and males. We first compared the average provisioning interval one day before and one day after the capture using two-sided paired t-tests to determine whether the birds returned to regular feeding behaviour after the initial capture. To determine if the average provisioning interval remained constant or if missed feeding time due to post-capture nest absence was later compensated for, we compared the average provisioning interval one and two days after the capture, again using a two-sided paired t-test for females and males separately.

We studied the longer-term implications of the protocol on fitness using first fledgling number and then fledgling mass and tarsus length as proxies. We first ran two generalised linear mixed-effect models with Poisson distribution fitting fledgling number (median = 5, range: 0-12) as the response variable and stress-induced corticosterone levels or provisioning latency as covariates in separate models, together with hatching date and clutch size. Nest ID and year were included as random factors. Next, we conducted four linear mixed-effect models with a Gaussian distribution fitting fledgling mass and tarsus length as response variables, respectively, and stress-induced corticosterone levels or provisioning latency as covariates. In each of these four models, we also included hatching date and clutch size as covariates. Since body mass can change throughout the day, we also added time of the day as a covariate for the two models looking at this variable. Sex and its interaction with either stress-induced corticosterone levels or provisioning latency were also included in the models. The random effects included in these models were parental bird ID, person ID (i.e., the person who took morphological data) and year.

Lastly, we examined whether the abandonment of a parent would lead to nest failure or reduce fledging success. For this, we fitted nest success (levels: no vs. yes) and fledgling number as response variables of two separate generalised linear mixed-effect models with Poisson distribution and a binomial distribution, respectively. For both models, we set the fixed factor return (levels: no vs. yes) in interaction with sex and added nest ID and year as random factors. In the model testing for fledgling number, we additionally included the hatch date of the offspring as a covariate to control for within-season differences.

### Ethical Note

All field procedures were conducted in compliance with the legal requirements of Germany and the EU and were approved by the cognizant regional governmental authority, the Regierungspräsidium von Oberbayern, Germany (license no. ROB-55.2-2532-Vet_02-13-204 & ROB-55.2-2532-Vet_02-15-25).

## 3. Results

Over the course of three breeding seasons, we captured and collected morphological data for 185 individual adult great tits. Of these 185 adults, 86 were captured repeatedly; 48 were recorded twice, 32 were recorded three times, five were recorded four times and one bird was even recorded five times. We also collected morphological data for 722 fledglings from 136 nests (1^st^ clutch: 83%, 2^nd^ clutch: 17%; N nests: 2020: 406, 2021: 185, 2022: 331).

We collected blood samples to quantify stress-induced corticosterone levels from 183 adult individuals, resulting in 299 samples. Nine samples had to be excluded due to their exceptionally high values (see 2.3 Corticosterone analysis), leaving us with 290 samples (2020: 81, 2021: 100, 2022: 109) from 181 individuals with 160 nests (1^st^ clutch: 83%, 2^nd^ clutch: 17%). Out of the 181 individuals, 48 were sampled twice, 26 were sampled three times and three were sampled four times. Average stress-induced sampling time was 28.70 ± 3.74 minutes (range: 21.00 - 40.67 min).

We collected behavioural data for 169 individuals. Out of these 169 individuals, 39 were recorded for two different breeding attempts, 33 were recorded for three attempts and eight were recorded for four attempts, resulting in 298 recordings (N females: 2020: 42, 2021: 55, 2022: 57; N males: 2020: 37, 2021: 55, 2022: 52) from 165 nests (1^st^ clutch: 82%, 2^nd^ clutch: 18%; N nests: 2020: 45, 2021: 61, 2022: 59). For 82 out of the 169 individuals, we had average provisioning interval data the two days before and after the capture. Out of these 82 individuals, 23 were recorded twice (N females: 12, N males: 11) and four individuals were recorded three times (N females: 2, N males: 2). This resulted in 113 paired recordings (N females: 59, N males: 54) of the average provisioning interval the two days before and the two days after the capture.

### 3.1 Short-term relationship of the ‘capture-handling-restraint protocol’ with physiology and behaviour

Baseline corticosterone levels (mean ± SD = 4.196 ± 2.633 ng/ml, range: 0.061-17.691 ng/ml) were lower compared to stress-induced corticosterone levels (mean ± SD = 28.687 ± 10.672 ng/ml, range: 3.950-66.479 ng/ml) upon capture (t(275) =-38.847, p-value < 0.000). These results hold after looking at each sex separately (females: t(138) =-27.125, p-value < 0.000; males: t(136) =-28.002, p-value < 0.000).

Out of 240 capture events, 217 captured birds returned to their nests (209 on the same day and eight on the following day) and 23 birds abandoned the nest (11 female individuals and 11 male individuals, with one male which abandoned twice, both times within one year). Parents resumed feeding on average after 2.53 ± 3.20 hours. Females and males had similar provisioning latencies of 2.47 ± 2.91 and 2.59 ± 3.52 hours, respectively. The provisioning latency differed significantly from the average provisioning interval the day before the capture for both sexes (females: t(58) =-6.158, p-value < 0.000; males: t(53) =-5.168, p-value < 0.000), which was 8.72 ± 3.57 minutes for females and 6.87 ± 1.97 minutes for males (females: min. = 4. 20 min, max. = 24.45 min; males: min. = 4.08 min, max. = 14.23 min; for more details see Supplementary Information 5).

While sampling time explained differences in physiology observed among male great tits, this was not the case for females. Handling time did not explain differences in observed behaviour for either sex. Sampling time tended to be negatively related to stress-induced corticosterone levels (Table 1A; Figure 2A) and handling time appeared to be negatively related to both abandonment rate (Table 2A) and provisioning latency (Table 1B; Figure 2B). However, of these relationships, only the relationship of sampling time and stress-induced corticosterone levels of male birds was significant (posterior probabilities for females and males: stress-induced corticosterone levels: > 74% and > 99%; abandonment: > 62% and > 52%; provisioning latency: > 72% and > 27%). Stress-induced corticosterone levels were positively correlated with provisioning latency, although this correlation too was not statistically significant (posterior probabilities: females > 13% and males > 56%; Figure 2; Table 1C).

**Figure 2.**
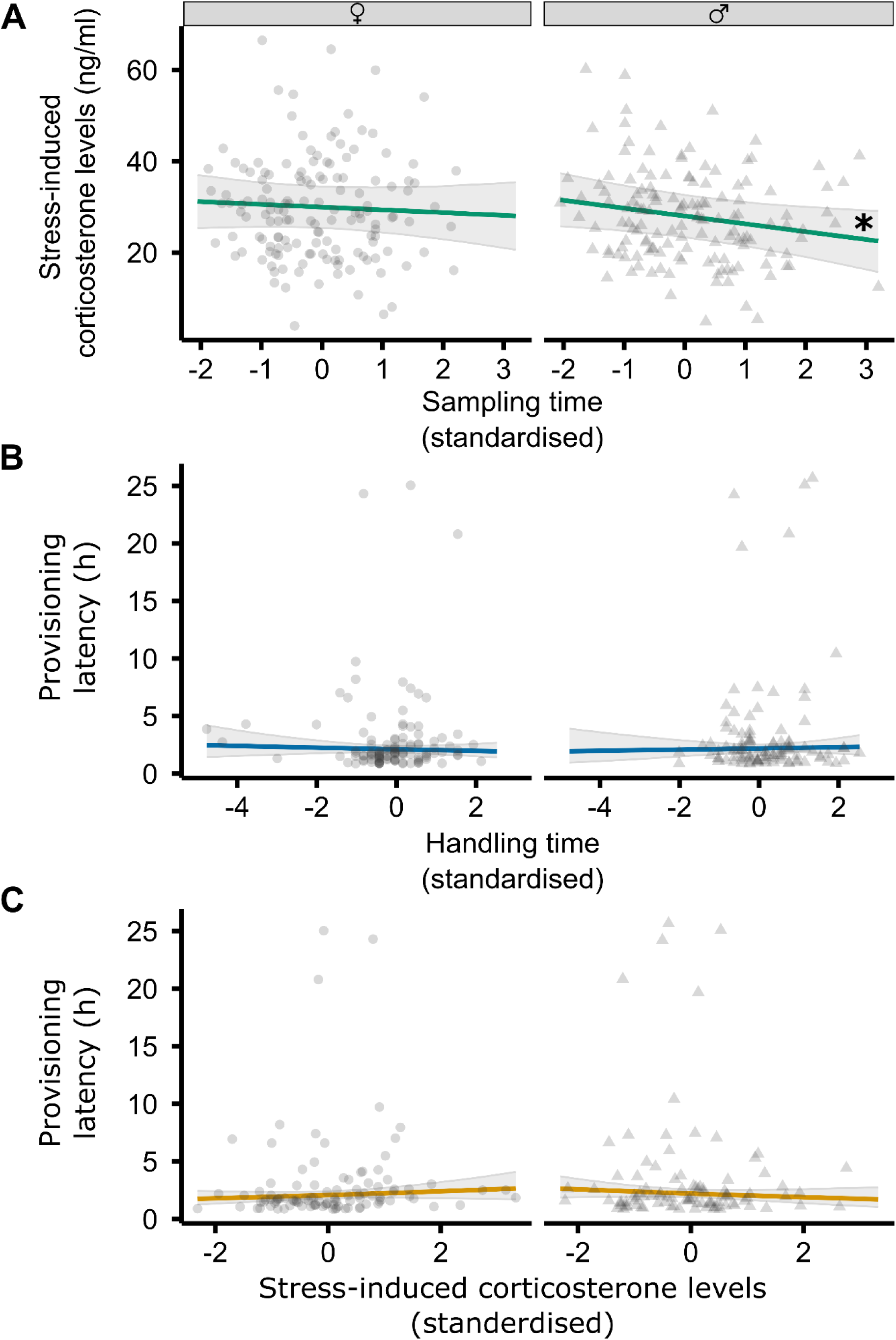
Relationship of sampling and handling time with physiology and behaviour for breeding female (circles) and male (triangles) wild great tits. Association of **A**) sampling time with stress-induced corticosterone levels, **B**) handling time with the provisioning latency (i.e., time between release and first return to the nest box) and **C**) stress-induced corticosterone levels with provisioning latency. Figures show mean estimates (coloured lines), 95% credible intervals (shaded grey) and raw data (grey symbols). The asterisk (*) indicates statistical significance for the relationship between sampling time and stress-induced corticosterone levels in males, meaning males that experienced longer sampling times had lower stress-induced corticosterone levels.

**Table 1.**
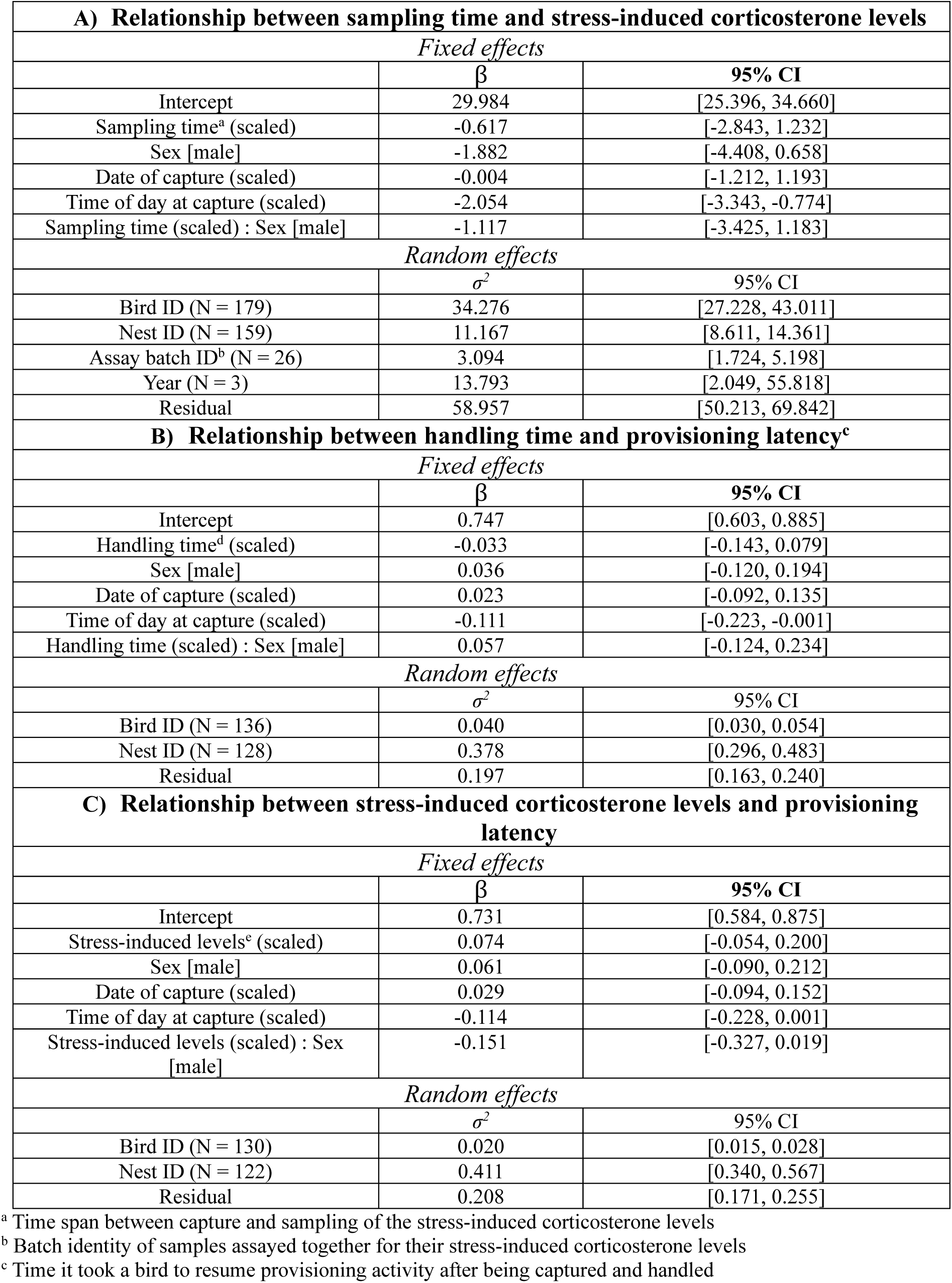
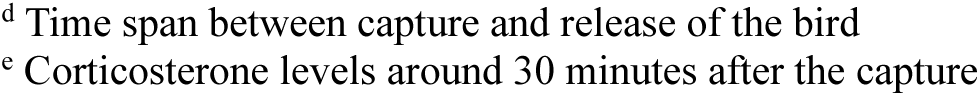
Results from mixed models testing the short-term relationships of sampling and handling protocol with physiology and behaviour. Statistical significance was considered when the CI did not overlap zero.

| **A) Relationship between sampling time and stress-induced corticosterone levels** |  |  |
| --- | --- | --- |
| *Fixed effects* |  |  |
| | $\beta$ | 95% CI |
| Intercept | 29.984 | [25.396, 34.660] |
| Sampling time<sup>a</sup> (scaled) | -0.617 | [-2.843, 1.232] |
| Sex [male] | -1.882 | [-4.408, 0.658] |
| Date of capture (scaled) | -0.004 | [-1.212, 1.193] |
| Time of day at capture (scaled) | -2.054 | [-3.343, -0.774] |
| Sampling time (scaled) : Sex [male] | -1.117 | [-3.425, 1.183] |
| *Random effects* |  |  |
| | $\sigma^2$ | 95% CI |
| Bird ID (N = 179) | 34.276 | [27.228, 43.011] |
| Nest ID (N = 159) | 11.167 | [8.611, 14.361] |
| Assay batch ID<sup>b</sup> (N = 26) | 3.094 | [1.724, 5.198] |
| Year (N = 3) | 13.793 | [2.049, 55.818] |
| Residual | 58.957 | [50.213, 69.842] |
| **B) Relationship between handling time and provisioning latency<sup>c</sup>** |  |  |
| *Fixed effects* |  |  |
| | $\beta$ | 95% CI |
| Intercept | 0.747 | [0.603, 0.885] |
| Handling time<sup>d</sup> (scaled) | -0.033 | [-0.143, 0.079] |
| Sex [male] | 0.036 | [-0.120, 0.194] |
| Date of capture (scaled) | 0.023 | [-0.092, 0.135] |
| Time of day at capture (scaled) | -0.111 | [-0.223, -0.001] |
| Handling time (scaled) : Sex [male] | 0.057 | [-0.124, 0.234] |
| *Random effects* |  |  |
| | $\sigma^2$ | 95% CI |
| Bird ID (N = 136) | 0.040 | [0.030, 0.054] |
| Nest ID (N = 128) | 0.378 | [0.296, 0.483] |
| Residual | 0.197 | [0.163, 0.240] |
| **C) Relationship between stress-induced corticosterone levels and provisioning latency** |  |  |
| *Fixed effects* |  |  |
| | $\beta$ | 95% CI |
| Intercept | 0.731 | [0.584, 0.875] |
| Stress-induced levels<sup>e</sup> (scaled) | 0.074 | [-0.054, 0.200] |
| Sex [male] | 0.061 | [-0.090, 0.212] |
| Date of capture (scaled) | 0.029 | [-0.094, 0.152] |
| Time of day at capture (scaled) | -0.114 | [-0.228, 0.001] |
| Stress-induced levels (scaled) : Sex [male] | -0.151 | [-0.327, 0.019] |
| *Random effects* |  |  |
| | $\sigma^2$ | 95% CI |
| Bird ID (N = 130) | 0.020 | [0.015, 0.028] |
| Nest ID (N = 122) | 0.411 | [0.340, 0.567] |
| Residual | 0.208 | [0.171, 0.255] |

<sup>a</sup> Time span between capture and sampling of the stress-induced corticosterone levels
<sup>b</sup> Batch identity of samples assayed together for their stress-induced corticosterone levels
<sup>c</sup> Time it took a bird to resume provisioning activity after being captured and handled
<sup>d</sup> Time span between capture and release of the bird <sup>e</sup> Corticosterone levels around 30 minutes after the capture

**Table 2.** Results from mixed models testing the relationship between abandonment and indicators of fitness. Statistical significance was considered when the CI did not overlap zero.

| **A) Relationship between handling time<sup>a</sup> and abandonment** |  |  |
| --- | --- | --- |
| *Fixed effects* |  |  |
| | $\beta$ | 95% CI |
| Intercept | 28.648 | [-67.162, 122.636] |
| Handling time | -27.543 | [-215.820, 136.776] |
| Sex [male] | -27.434 | [-149.576, 91.168] |
| Handling time: Sex [male] | 50.091 | [-149.559, 255.413] |
| *Random effects* |  |  |
| | $\sigma^2$ | 95% CI |
| Nest ID (N = 142) | 1247.662 | [967.369, 1577.736] |
| Year (N = 3) | 0.011 | [0.000, 0.057] |
| **B) Relationship between abandonment and nest success<sup>b</sup>** |  |  |
| *Fixed effects* |  |  |
| | $\beta$ | 95% CI |
| Intercept | 10.586 | [-31.104, 53.190] |
| Return [yes] | 4.195 | [-43.832, 51.905] |
| Sex [male] | 0.976 | [-47.817, 48.473] |
| Return [yes] : Sex [male] | -0.971 | [-51.857, 50.658] |
| *Random effects* |  |  |
| | $\sigma^2$ | 95% CI |
| Nest ID (N = 142) | 2311.270 | [1766.692, 2929.550] |
| Year (N = 3) | 25.025 | [0.909, 128.759] |
| **C) Relationship between abandonment and fledgling number** |  |  |
| *Fixed effects* |  |  |
| | $\beta$ | 95% CI |
| Intercept | 2.128 | [0.611, 3.640] |
| Return [yes] | 0.706 | [0.234, 1.182] |
| Sex [male] | 0.306 | [-0.240, 0.859] |
| Hatch date (numeric) | -0.261 | [-0.534, 0.013] |
| Return [yes] : Sex [male] | -0.305 | [-0.867, 0.263] |
| *Random effects* |  |  |
| | $\sigma^2$ | 95% CI |
| Nest ID (N = 142) | 0.140 | [0.112, 0.171] |
| Year (N = 3) | 0.079 | [0.007, 0.247] |

<sup>a</sup> Time span between capture and release of the bird
<sup>b</sup> Binomial variable; 'nest success' (any number of fledged offspring) or 'nest failure' (no fledged offspring)

### 3.2 Relationship of ambient temperature, parental condition and brood quality with behaviour

Ambient temperature at capture (Table 3A), parental condition (Tables 3B & 3C) and brood quality (Tables 3D & 3E) did not explain variation in provisioning latency. Great tits captured and handled during warmer times of the day seemed to return to their nests later than those captured during cooler periods. However, these relationships were not statistically significant (posterior probabilities for females and males: > 18% and > 42%). While females appeared to take longer to return to their nests when caring for a second clutch or for more chicks, males returned more quickly. Both sexes seemed to return faster when nestlings were older. However, once again, none of these relationships were statistically significant (posterior probabilities for females and males: clutch number: > 4% and > 91%; number of nestlings: > 40% and > 93%; nestling age: > 59% and > 67%).

**Table 3.** Results from mixed models testing the relationship between non-handling related variables with provisioning latency. Statistical significance was considered when the CI did not overlap zero.

| **A) Relationship between temperature at capture and provisioning latency** |  |  |
| --- | --- | --- |
| *Fixed effects* |  |  |
| | $\beta$ | 95% CI |
| Intercept | 0.755 | [0.613, 0.899] |
| Temperature at capture (°C) | 0.067 | [-0.076, 0.211] |
| Sex [male] | 0.041 | [-0.115, 0.202] |
| Capture date (scaled) | 0.007 | [-0.128, 0.139] |
| Capture time (scaled) | -0.125 | [-0.241, -0.005] |
| Temperature at capture (scaled) : Sex [male] | -0.084 | [-0.228, 0.057] |
| *Random effects* |  |  |
| | $\sigma^2$ | 95% CI |
| Bird ID (N = 135) | 0.051 | [0.038, 0.069] |
| Nest ID (N = 126) | 0.377 | [0.296, 0.481] |
| Residual | 0.188 | [0.155, 0.230] |
| **B) Relationship between parental condition and provisioning latency (females)** |  |  |
| *Fixed effects* |  |  |
| | $\beta$ | 95% CI |
| Intercept | -0.247 | [-5.439, 4.885] |
| Mass (g) | -0.114 | [-0.281, 0.052] |
| Tarsus length (mm) | 0.154 | [-0.100, 0.412] |
| Age [adult] | -0.091 | [-0.361, 0.168] |
| Parasites [yes] | 0.103 | [0.145, -0.346] |
| *Random effects* |  |  |
| | $\sigma^2$ | 95% CI |
| Bird ID (N = 75) | 0.273 | [0.206, 0.358] |
| Year (N = 3) | 0.004 | [0.000, 0.021] |
| Residual | 0.227 | [0.176, 0.300] |
| **C) Relationship between parental condition and provisioning latency (males)** |  |  |
| *Fixed effects* |  |  |
| | $\beta$ | 95% CI |
| Intercept | 0.257 | [-3.828, 4.372] |
| Mass (g) | -0.094 | [-0.300, 0.110] |
| Tarsus length (mm) | 0.121 | [-0.138, 0.375] |
| Age [adult] | -0.348 | [-0.725, 0.025] |
| Parasites [yes] | 0.207 | [-0.113, 0.532] |
| **D) Relationship between brood quality and provisioning latency (females)** |  |  |
| *Fixed effects* |  |  |
| | $\beta$ | 95% CI |
| Intercept | 0.780 | [-0.557, 2.129] |
| Clutch number [2] | 0.275 | [-0.028, 0.578] |
| Age nestlings (days) | -0.015 | [-0.148, 0.118] |
| Clutch size | 0.011 | [-0.078, 0.101] |
| *Random effects* |  |  |
| | $\sigma^2$ | 95% CI |
| Parental bird ID (N = 74) | 0.279 | [0.210, 0.368] |
| Year (N = 3) | 0.001 | [0.000, 0.006] |
| Residual | 0.251 | [0.195, 0.332] |
| **E) Relationship between brood quality and provisioning latency (males)** |  |  |
| *Fixed effects* |  |  |
| | $\beta$ | 95% CI |
| Intercept | 1.730 | [0.018, 3.414] |
| Clutch number [2] | -0.315 | [-0.780, 0.147] |
| Age nestlings (days) | -0.037 | [-0.204, 0.132] |
| Clutch size | -0.096 | [-0.225, 0.031] |

| *Random effects* |  |  |
| --- | --- | --- |
| | $\sigma^2$ | 95% CI |
| Parental bird ID (N = 61) | 0.073 | [0.047, 0.111] |
| Residual | 0.561 | [0.429, 0.754] |

### 3.3 Longer-term relationship of the ‘capture-handling-restraint protocol’ with proxies for reproductive success

Parents that returned to their nests after capture and handling resumed provisioning with intervals on the following day comparable to the day preceding capture (females and males: N = 59, 54; average provisioning interval on the following day after capture = 8.09 ± 5.43 min, 6.91 ± 2.47 min; p-values = 0.229, 0.856; Supplementary Table 1). Stress-induced corticosterone levels appeared positively associated with fledgling number and negatively associated with fledgling mass and tarsus length. But these relationships were not statistically significant (posterior probabilities for females and males: fledgling number: > 44% and > 54%; mass: > 89% and > 12%; tarsus length only for males: > 23%). However, females with lower stress-induced corticosterone levels raised nestlings with significantly longer tarsi (posterior probability > 99%; Figure 3; Table 4C). Provisioning latency seemed positively related to all three fitness proxies, but none of these relationships were statistically significant (posterior probabilities for females and males: fledgling number: > 26% and > 50% (Table 5A); fledgling mass: > 23% and > 57% (Table 5B); fledgling tarsus length: > 41% and 47% (Table 5C); Figure 3). The return of a parent did not explain whether a nest was successful or not (posterior probabilities for both sexes > 43%; Table 2B). However, individuals that returned to the nest raised more fledglings than individuals that did not (posterior probabilities for both sexes > 0.1%; Table 2C).

**Figure 3.**
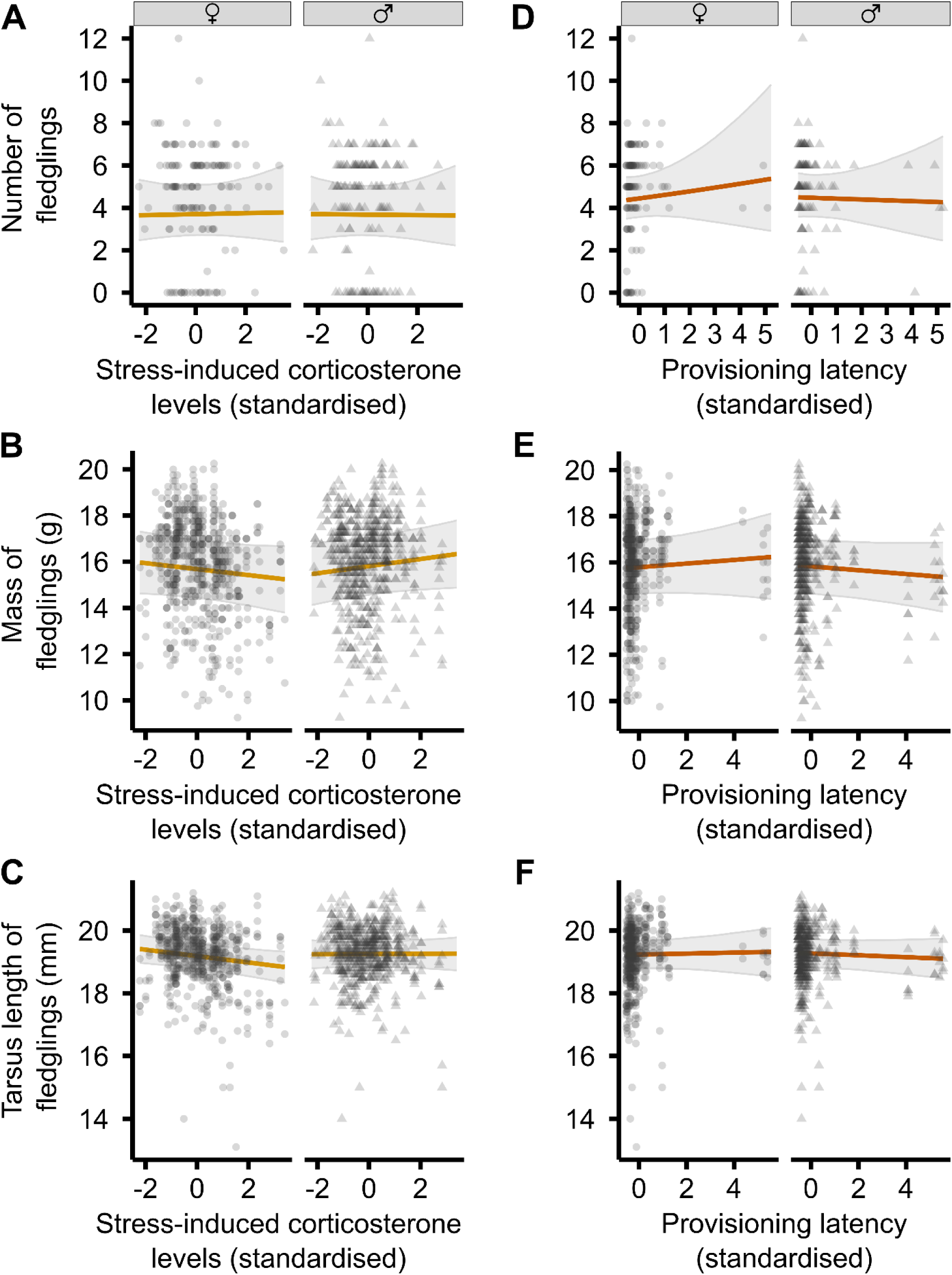
Relationship of physiology and behaviour with fitness indicators for breeding female (circles) and male (triangles) wild great tits. Association of stress-induced corticosterone levels with **A**) number of fledglings, **B**) mass of fledglings and **C**) tarsus length of fledglings. Association of provisioning latency (i.e., time between release and first return to the nest box) with **D**) number of fledglings, **E**) mass of fledglings and **F**) tarsus length of fledglings. Figures show mean estimates (coloured lines: yellow: stress-induced corticosterone levels, orange: provisioning latency), 95% credible intervals (shaded grey) and raw data (grey symbols).

**Table 4.** Results from mixed models testing the relationship between the physiology of breeding great tits with indicators of fitness. Statistical significance was considered when the CI did not overlap zero.

| **A) Relationship between stress-induced corticosterone levels<sup>a</sup> and fledgling number** |  |  |
| --- | --- | --- |
| *Fixed effects* |  |  |
| | $\beta$ | 95% CI |
| Intercept | 1.310 | [0.998, 1.624] |
| Stress-induced levels (scaled) | 0.007 | [-0.089, 0.106] |
| Sex [male] | -0.006 | [-0.122, 0.112] |
| Hatch date (scaled) | -0.076 | [-0.188, 0.034] |
| Clutch size | 0.075 | [-0.035, 0.183] |
| Stress-induced levels (scaled) : Sex [male] | -0.010 | [-0.144, 0.126] |
| *Random effects* |  |  |
| | $\sigma^2$ | 95% CI |
| Nest ID (N = 157) | 0.309 | [0.257, 0.366] |
| Year (N = 3) | 0.092 | [0.012, 0.261] |
| **B) Relationship between stress-induced corticosterone levels and fledgling mass** |  |  |
| *Fixed effects* |  |  |
| | $\beta$ | 95% CI |
| Intercept | 16.875 | [15.388, 18.335] |
| Stress-induced levels (scaled) | -0.127 | [-0.327, 0.079] |
| Sex [male] | 0.127 | [-0.281, 0.540] |
| Hatch date (scaled) | 0.102 | [-0.022, 0.223] |
| Clutch size | -0.164 | [-0.265, -0.058] |
| Capture time of the chicks (scaled) | 0.138 | [-0.009, 0.285] |
| Stress-induced levels (scaled) : Sex [male] | 0.289 | [-0.002, 0.580] |
| *Random effects* |  |  |
| | $\sigma^2$ | 95% CI |
| Parental bird ID (N = 161) | 1.405 | [1.212, 1.621] |
| Observer ID (N = 5) | 0.240 | [0.080, 0.498] |
| Year (N = 3) | 0.984 | [0.616, 1.453] |
| Residual | 2.007 | [1.855, 2.173] |
| **C) Relationship between stress-induced corticosterone levels and fledgling tarsus length** |  |  |
| *Fixed effects* |  |  |
| | $\beta$ | 95% CI |
| Intercept | 18.846 | [18.326, 19.361] |
| Stress-induced levels (scaled) | -0.103 | [-0.187, -0.018] |
| Sex [male] | 0.066 | [-0.084, 0.217] |
| Hatch date (scaled) | -0.023 | [-0.079, 0.035] |
| Clutch size | 0.046 | [0.001, 0.091] |
| Stress-induced levels (scaled) : Sex [male] | 0.107 | [-0.022, 0.235] |
| *Random effects* |  |  |
| | $\sigma^2$ | 95% CI |
| Parental bird ID (N = 161) | 0.143 | [0.118, 0.171] |
| Observer ID (N = 5) | 0.066 | [0.023, 0.140] |
| Year (N = 3) | 0.067 | [0.029, 0.124] |
| Residual | 0.564 | [0.522, 0.610] |

<sup>a</sup> Corticosterone levels around 30 minutes after capture

**Table 5.** Results from mixed models testing the relationship between behaviour of breeding great tits and indicators of their fitness. Statistical significance was considered when the CI did not overlap zero.

| **A) Relationship between provisioning latency<sup>a</sup> and fledgling number** |  |  |
| --- | --- | --- |
| *Fixed effects* |  |  |
| | $\beta$ | 95% CI |
| Intercept | 1.491 | [1.271, 1.704] |
| Provisioning latency (scaled) | 0.036 | [-0.073, 0.144] |
| Sex [male] | 0.008 | [-0.117, 0.135] |
| Hatch date (scaled) | -0.073 | [-0.161, 0.014] |
| Clutch size | 0.097 | [0.013, 0.179] |
| Provisioning latency (scaled) : Sex [male] | -0.045 | [-0.179, 0.089] |
| *Random effects* |  |  |
| | $\sigma^2$ | 95% CI |
| Nest ID (N = 127) | 0.082 | [0.064, 0.102] |
| Year (N = 3) | 0.043 | [0.003, 0.149] |
| **B) Relationship between provisioning latency and fledgling mass** |  |  |
| *Fixed effects* |  |  |
| | $\beta$ | 95% CI |
| Intercept | 15.785 | [14.609, 16.948] |
| Provisioning latency (scaled) | 0.101 | [-0.168, 0.365] |
| Sex [male] | 0.032 | [-0.402, 0.472] |
| Hatch date (scaled) | 0.054 | [-0.077, 0.185] |
| Clutch size | -0.188 | [-0.333, -0.045] |
| Capture time of the chicks (scaled) | -0.075 | [-0.233, 0.082] |
| Provisioning latency (scaled) : Sex [male] | -0.175 | [-0.495, 0.139] |
| *Random effects* |  |  |
| | $\sigma^2$ | 95% CI |
| Parental bird ID (N = 131) | 1.302 | [1.100, 1.536] |
| Observer ID (N = 5) | 0.810 | [0.258, 1.781] |
| Year (N = 3) | 0.499 | [0.105, 1.461] |
| Residual | 2.125 | [1.951, 2.323] |
| **C) Relationship between provisioning latency and fledgling tarsus length** |  |  |
| *Fixed effects* |  |  |
| | $\beta$ | 95% CI |
| Intercept | 19.235 | [18.796, 19.671] |
| Provisioning latency (scaled) | 0.013 | [-0.098, 0.124] |
| Sex [male] | 0.037 | [-0.136, 0.216] |
| Hatch date (scaled) | -0.030 | [-0.093, 0.033] |
| Clutch size | 0.086 | [0.018, 0.153] |
| Provisioning latency (scaled) : Sex [male] | -0.044 | [-0.180, 0.095] |
| *Random effects* |  |  |
| | $\sigma^2$ | 95% CI |
| Parental bird ID (N = 131) | 0.167 | [0.136, 0.201] |
| Observer ID (N = 5) | 0.156 | [0.041, 0.362] |
| Year (N = 3) | 0.040 | [0.004, 0.127] |
| Residual | 0.606 | [0.556, 0.662] |

<sup>a</sup> Time it took a bird to resume provisioning activity after being captured and handled

## 4. Discussion

We examined the short-and longer-term effects of a standardised ‘capture-handling-restraint protocol’ by analysing its relationship behavioural and physiological responses of adult great tits during the nestling provisioning phase and the relationship of these responses with reproductive success. Overall, in 90.4% of capture events, birds returned to their nests. The protocol strongly affected adult physiological and behavioural traits on the short term, but had no longer-term consequences on behaviour. Moreover, except for abandonment, these short-term physiological and behavioural changes did not explain variation in fledgling condition and number, likely because, once parents returned to their nests, they resumed normal average provisioning intervals. While abandonment reduced the number of fledglings, it did not affect overall nest success or failure.

On a short-time scale, we found that fewer than 10% of capturing events (23 out of 240) resulted in parents abandoning their nests after being captured and this was not related to the duration of handling. Nonetheless, it is worth noting that the abandonment by one parent did not automatically lead to a complete nest failure, likely due to continued provisioning by the other parent. This suggests that, if a parent abandons the nest when the nestlings are between seven and twelve days old, the other parent may be capable of caring for the nestlings until fledging, albeit less efficiently, as indicated by the decreased number of fledglings in nests where one parent abandoned We also found significant increases in corticosterone levels following capture, confirming the expected physiological activation of the HPA axis. However, the lack of a relationship between sampling time and stress-induced corticosterone levels in females and the negative correlation for males did not align with our expectations. The females’ result might be explained by the fact that most bird species reach their peak corticosterone levels in response to the initial encounter of an unpredictable stimulus at around 30 min (Baugh et al., 2012; Monaghan and Spencer, 2014; Romero et al., 2000; Wilson and McMahon, 2006; Wingfield et al., 1998), which is approximately when we sampled most great tits (i.e., 28.70 min). Consistently, Huber et al. (2021) found a difference in stress-induced concentrations between 15 and 30 min, irrespective of the three different handling protocols used. One possible explanation for our findings in males is that there are sex differences in HPA axis regulation during the breeding season, with the negative feedback being induced earlier in males than in females. However, to our knowledge, there are no studies in birds supporting this hypothesis. We only found results indicating the opposite: breeding males being insensitive to negative glucocorticoid feedback (Astheimer et al., 1994). Furthermore, free-living animals’ stress responses are context-dependent and modulated by various other factors (e.g., season and weather, development, previous experiences or remaining reproductive opportunities), making it difficult to connect the consequences to a single immediate stressor (reviewed by Bonier et al., 2009; Crespi et al., 2013; Kitaysky et al., 1999; Romero et al., 2000; Wingfield et al., 1998). For example, Lynn et al. (2010) studied whether one capture experience can influence the stress response to the same protocol later on. They subjected free-living eastern female bluebirds to a capture-handling-restraint protocol, distinguishing between three test groups: birds that were captured twice (once during their first and once during their second brood, i.e., repeaters) or birds that were subjected to the protocol only once (either during their first or second brood). They found a decline in corticosterone secretion in repeaters compared to birds experiencing the protocol for the first time, indicating that they acclimated or habituated to the protocol after only one experience. This suggests that a singular previous stressful experience can significantly influence HPA responsiveness.

Likewise, to the significant increases in corticosterone levels following capture, provisioning latency was also significantly delayed, suggesting a behavioural stress response. But its magnitude was not related to the duration of handling. Our result aligns with the results of a study on blue tits, where researchers retained birds on average for 30 minutes (range 7-84 min; Schlicht & Kempenaers, 2015). It is possible that the birds displayed a behavioural response immediately after capture and/or that the response was related to other factors not measured in our study.

Stress-induced corticosterone levels also did not explain the duration birds stayed away from their nests. This result is in line with Lane et al. (2025), who reported that neither baseline nor stress-induced corticosterone concentrations were correlated with return latency in female song sparrows, arguing that corticosterone levels are not the only mechanism that mediates behavioural stress responses. To test if an increase in stress-induced corticosterone levels affects provisioning latency following capture and handling, one could conduct a follow-up study and experimentally manipulate corticosterone’s actions (see review by Hau et al., 2016). In general, experimental studies have linked glucocorticoids to parental provisioning in birds. For instance, administration of corticosterone implants to pied flycatcher parents reduced provisioning intervals and, at high concentrations, induced brood abandonment (Silverin, 1986). Likewise, corticosterone implants in male, white-crowned sparrows (*Zonotrichia albicollis*) from the high-provisioning (tan-striped) morph reduced offspring feeding rates, while the glucocorticoid receptor antagonist RU486 induced an increase in provisioning in the low-provisioning, white-striped morph (Almasi et al., 2008; Horton and Holberton, 2009). Also, daily pulses of high corticosterone exposure in incubating female tree swallows (*Tachycineta bicolor*) reduced later provisioning rates, especially when caring for large broods (Vitousek et al., 2018). Yet, these results concern provisioning rates and not provisioning latency after an acute stressor and therefore might represent different underlying mechanisms. Taken together, it is likely that the increase in corticosterone helped great tit parents in our study cope with the unexpected experience of being captured and supported changes in their provisioning behaviour (e.g., replenishing energy before returning to the nest). However, differences in provisioning latency are likely driven by factors other than stress-induced corticosterone levels (and also not by ambient temperature, parental condition or brood quality). For example, variation in the timing of behavioural resumption may be associated with the efficiency of HPA axis downregulation (Hau et al., 2016; Lattin and Kelly, 2020; Romero, 2004; Taff et al., 2018). Experimental work in eastern bluebirds has shown that more effective negative feedback mechanisms might predict an individual’s resilience in response to transient stressors (Taff et al., 2018). A promising avenue for future research would therefore be to relate provisioning latency to glucocorticoid negative feedback. Likewise, to investigate physiological mechanisms mediating such behaviour, one could focus on both corticosterone and prolactin, the latter known as the ‘parental hormone’, since these hormones influence parental behaviour and may affect different components of the stress response (reviewed by Angelier & Chastel, 2009; Chastel et al., 2005; Ouyang et al., 2013). Additionally, one could explore the social dynamics within breeding pairs, as in great tits, the behaviour of one parent might influence the other (Székely et al., 1996), as well as genetic differences in the behavioural stress response (Mutzel et al., 2013; Van Oers et al., 2004).

On a longer-time scale, great tit parents that returned to their nests resumed normal provisioning behaviour. This result is in line with Schlicht & Kempenaers (2015), who found similar time spans between nest blue tit visits preceding capture and after the initial return. Contrary to our predictions, neither stress-induced corticosterone levels nor provisioning latency explained variation in fitness proxies, except for fledglings of parents with higher corticosterone levels having shorter tarsus lengths. Given that tarsus length is a morphological trait largely determined by genetics (Husby et al., 2011; Noordwijk et al., 1988) and that birds are exposed to multiple challenging events during the breeding season that may trigger a stress response (Huber et al., 2021; Lack, 1968; Soulsbury et al., 2020), it seems unlikely that a single capture event caused nestlings to develop shorter tarsi. This raises the question: why does an interruption of several hours in a usually frequent parental care behaviour have no major impact on reproductive success? Was the missing food delivery subsequently compensated for? Curiously, we did not find a significant increase in provisioning that would support the idea of compensation after a parent’s return to the nest. Despite clear physiological and behavioural responses to the ‘capture-handling-restraint protocol’, the absence of measurable consequences for offspring number or condition, aside from a weak relationship with tarsus length, suggests a large degree of resilience in this (and many other) species during the provisioning period. These findings, therefore, contribute to a growing body of literature that questions the longer-term impacts of acute, short-term stressors on reproductive success in wild bird populations (Lynn et al., 2010; Nicolaus et al., 2008; Ouyang et al., 2013; Schlicht and Kempenaers, 2015). However, this should not be interpreted as evidence that such disturbances are inconsequential. Our study naturally lacked a true control group of uncaptured individuals (because assessing corticosterone levels requires capture and handling), limiting our ability to detect subtle differences in longer-term relationships with reproductive output.

## 5. Conclusion

Capture, handling and sampling are integral to research on wild animals in fields of biology as diverse as ecology, evolution, physiology, genetics and conservation. Yet our understanding of the short-and long-term implications of these procedures has remained limited. In this study, we assessed immediate physiological and behavioural responses to a standardised ‘capture-handling-restraint protocol’, as well as longer-term behaviour and reproductive success over three consecutive breeding seasons. While the protocol triggered clear acute stress responses, our findings suggest that great tits are behaviourally resilient, with no detectable long-term implications on reproductive success. However, these conclusions must be interpreted with caution. The absence of an uncaptured control group limits our ability to fully assess potential subtle or cumulative relationships. It also remains possible that such brief disruptions may have longer-term consequences not captured by our metrics or that become evident only across successive breeding attempts. To advance both scientific understanding and ethical standards, future research should prioritise the inclusion of true control groups, consider the role of social dynamics within breeding pairs and monitor reproductive success over multiple breeding seasons. Additionally, comparative studies would be valuable for determining species’ sensitivity to capture-handling protocols. These studies should include species with divergent life-history strategies, varying reproductive investment per breeding bout and different survival rates. Such efforts are essential for refining best practices in field research, ensuring that the burden on the welfare of the wild animals studied does not outweigh the benefits of scientific data collection. This approach will enable researchers and regulatory agencies responsible for evaluating experimental studies on wild populations to conduct and assess scientific research responsibly and ethically.

## Acknowledgements

We would like to dedicate this paper to the memory of Michaela (Ela) Hau, whose insight, dedication and passion for research greatly shaped this study. Her contributions and enthusiasm continue to inspire us.

We highly appreciate the help from Tanguy Deville, Carlotta Bonaldi, Michi Stiegler and Natalia Perez with field and laboratory work. Many thanks to the Erzdiozöse München und Freising, especially K. Meindl, B. Vollmar and M. Laußer for allowing us to work in their forest. The Hau-Goymann laboratory group provided input that greatly improved data collection and manuscript preparation.

## Authors’ contributions

Conceptualisation: all authors. Data curation: FF, LM and CD. Formal analysis: FF and LM. Funding acquisition: MH. Investigation: all authors. Methodology: all authors. Project administration: LM and MH. Resources: MH and CD. Software: FF, LM and CD. Supervision: LM and MH. Validation: FF, LM and CD. Visualisation: FF and LM. Writing - original draft: FF and LM. Writing - review and editing: all authors.

## Data availability statement

All data and code are available at Zenodo: https://zenodo.org/records/19094415?token=eyJhbGciOiJIUzUxMiJ9.eyJpZCI6ImMxNDlhY TRlLWYwYmMtNGQ2MS05ZTczLTNmOGM3NmE4ZDI5OSIsImRhdGEiOnt9LCJyYW5k b20iOiJlZWZiMGZiMmQwMTUyZDQ3NWViNTJkMDVjMDA3M2FlNSJ9.kAil33Tx3Qw 3uBx-ZINQ7Pz52LTImKv3OaHPeWvUmjXk7pk_FVeETN4LV9XHz3R2m7jh77kbXtfbM2xftdhT 8A

## Declaration of Generative AI and AI-assisted technologies in the writing process

During the preparation of this work, the authors used ChatGPT in order to improve readability and language. After using this tool, the authors reviewed and edited the content as needed and take full responsibility for the content of the publication.

