## Supplementary Information for "A ‘capture-handling-restraint protocol’ elicits short-term stress responses in wild great tits (*Parus major*) but has little impact on reproductive success" (from another version of this preprint: version 1, 2026-03-19)

### Supplementary Information 1: Corticosterone Control Samples

We included control samples in each extraction batch that underwent the same extraction and analysis procedure as the samples. These control samples were analysed on the same assay plate as the samples from the respective batch. We included two positive quality control samples (QC 1+2) and one negative control sample in each batch. In total, we extracted and analysed 26 batches and plates, respectively.

The negative control sample contained only ultrapure water and was at the first and last position of each extraction batch. These controls were only measured as single measurements, as we only wanted to detect potential contamination with these controls. All negative control samples were above 90% binding. The detection limit of each plate was calculated as the mean optical density of the maximum binding (Zero, B0) duplicate wells minus two times the standard deviation between the duplicates. The mean detection limit across all assay plates was at 98.2 +/- 2.6 per cent binding, corresponding to a mean corticosterone level of 3.8 +/- 5.7 pg/ml. All samples used in the analysis were measured above the detection limit of the respective plate on which they were measured. Two positive control samples were measured in duplicate twice on each plate and were used to calculate intra- and inter-assay variation. The two positive control samples consisted of stripped chicken plasma spiked with Arbor Assays corticosterone stock solution (100,000 pg/ml corticosterone, Catalogue Number C151).

Control sample QC 1 had an absolute corticosterone level of 5,550 pg/ml and was measured at a 1:30 dilution factor. This resulted in a measurement at around 62% binding and an average corticosterone level of 199.3 +/- 29.3 pg/ml. Most baseline samples were measured in this range of the assay with dilution factors between 1:10-1:15, depending on available plasma sample volume. Control sample QC 2 had an absolute corticosterone level of 14,700 pg/ml and was measured at a 1:19 dilution factor. This resulted in a measurement at around 35% binding and an average level of 796.3 +/- 69.3 pg/ml. Stress-induced samples were measured in this range of the assay with dilution factors of 1:40-1:120, depending on available plasma sample volume.

Intra-assay variation was calculated by averaging the coefficient of variation between duplicate measurements of the control samples across all plates. Intra-assay variation was 13 ± 10% for QC 1, and 8.1 ± 4.5% for QC 2. Inter-assay variation was calculated by calculating the coefficient of variation across the corticosterone concentrations of the two control samples across plates. Inter-assay variation was 15.9% for QC1, and 9.1% for QC. Note that these values include not just variation on assay performance across plates, but also variation in extraction efficiency.

### Supplementary Information 2: Validation of the Automated Feeding Behaviour

Validating RFID data through visual observations, such as video recordings, is crucial for ensuring accurate behavioural estimates and accurately reflecting results. The reliability and consistency of RFID devices can vary significantly between studies, making it essential to perform validations within each specific study rather than relying on validations from other research. Unfortunately, many published studies fail to report the methodologies used for validating data accuracy and there is currently no standardised approach for RFID data validation (Hughes et al., 2021; Iserbyt et al., 2018; Nomano et al., 2014). Therefore, we performed a video validation in 2020, where we compared observations from the videos with the registered RFID data to measure the accuracy of automated feeding recordings. The videos were recorded between 0700 and 1300 hours when the nestlings were 7 or 12 days old to have measurements of the regular behaviour preceding and following the capture. The analysed duration was an hour per video, but the analysis only started after the video had already been running for at least 30 minutes, to ensure the birds were not behaving differently due to human disturbance. Our validation sample size consisted of 26 individuals from 13 different nests, of which one nest was observed twice, on day 7 and day 12, leading to 7 nest recordings on day 7 and 7 nest recordings on day 12. The results showed that the entries were registered reliably, with a 94.3% total registry rate. Females tended to be recorded more reliably (98.0% registry rate) than males (89.2% registry rate). However, the exits were only registered partially (22.4% registry rate) with little variation between sexes (females: 21.2% registry rate; males: 24.2% registry rate) but a high variation between individuals (i.e., some birds were not registered at all and for some birds every exit was recorded). This is probably due to the high speed at which birds are exiting, so the PIT tag could not be activated and read by the reader in time.

#### Supplementary Information 3: Data Curation of the Automated Feeding Behaviour

Although the RFID system is designed to detect the direction of movement, it frequently failed to do so reliably and often recorded the same movement twice. To have one record for each time a bird entered its nest box and avoid including duplicate reads or exits as entries, we applied a “cut-off rule” and excluded those recordings that occurred within 2 minutes. This approach is based on the previously performed validation and follows similar time frames found in other studies (e.g., Iserbyt et al., 2018; Nomano et al., 2014; Wilkin et al., 2009). It ensures that at least 90% of the time spent in the nest is captured (90th percentile of all birds: 1.87 min), while providing a safe buffer to distinguish between individual feeding events, as the average interval between consecutive feedings is 4.28 min.

### Supplementary Information 4: Validation and data curation of Latency

We performed a second validation in 2022, following the same protocol as previously described (Supplementary Information 2). We video recorded great tits after the birds were captured and subsequently released, with varying durations (mean = 2.5 h, min. = 0.8 h; max. = 5.7 h), depending on the scheduling constraints of fieldwork. We recorded videos for 15 nests post-capture, covering both parents in 13 nests and only the females in 2 nests due to the unavailability of the male (N = 28). Among the nests with both parents captured, 9 were caught on the same day, while the remaining ones were captured over two consecutive days. The analysis of the videos unveiled that the entry at the time of release was documented for only 15 out of the 28 individuals, resulting in a 46% rate of unregistered entries. One probable cause for this lack of registration could be attributed to the method of placing the birds back into the nest box. The positioning of the legs differs when the birds are put through the entrance hole by hand, in contrast to a bird entering on its own. This variance in leg positioning likely alters the angle of the PIT tag to the RFID reader, making it potentially more challenging for the reading device to register the entry (reviewed by Bonter & Bridge, 2011). Another likely explanation for the missing records is that the birds pass the reader too quickly when they enter the nest box immediately after being released (Hughes et al., 2021; Iserbyt et al., 2018). However, the validation showed that, for birds whose entry at release was manually recorded, the average difference from the release time noted in the field was just one minute. This supports the use of the release time from the field notes without significantly affecting the measurement of provisioning latency. Also, contrary to the previous validation, exits were recorded more frequently than when the birds exhibited normal behaviour. Exits were documented in 12 out of the 28 individuals, resulting in a 42.9% rate of registered exits. This is likely attributable to a more cautious behaviour when the birds exited the nest box again, which could also be observed in videos. Furthermore, the time spent inside the nest box after being released (mean = 15 min 44 s, min. = 4 min 59 s, max. = 40 min 25 s) was considerably longer than the average duration of a nest visit recorded during the monitoring of normal feeding behaviour (mean = 47 s, min. = 0 min 0 s, max. = 14 min 20 s). This behaviour could be attributed to their need for recovery or an attempt to avoid encountering additional predators.

This means that, based on our initial assumption from the first validation (that the exit would typically not be recorded and the next movement registered by the RFID reader after release would indicate the bird's return to the nest), in over 40% of cases, we would have incorrectly identified the exit as the return movement. The video analysis revealed that no bird returned in less than one hour after being released, rendering the substantial number of entries registered within the initial 20 minutes and even within the initial hour, highly improbable. Consequently, we disregarded recorded exits during the data analysis phase. We estimated an average provisioning latency as a preliminary insight into what to anticipate in the actual analyses; a mean of 3.88 h. Moreover, the validation demonstrated that every bird that returned during the recordings was carrying food, leading us to infer that the return of the parent was indeed associated with provisioning.

### Supplementary Information 5: Results

**Table 1.** Mean, standard deviation, minimum and maximum of variables indicating provisioning behaviour regarding the capture, first for all parents and then for females and males separately.

|  | Average provisioning interval two days prior to the capture  [min] | | Average provisioning interval one day prior to the capture  [min] | | Provisioning latency after the capture  [h] | | Average provisioning interval one day after the capture  [min] | | Average provisioning interval two days after the capture  [min] | |
| --- | --- | --- | --- | --- | --- | --- | --- | --- | --- | --- |
| **All parents (N=113)** | | | | | | | | | | |
| Mean | 8.43 | | 8.43 | | 2.53 | | 7.53 | | 7.89 | |
| Standard deviation | 3.87 | | 3.05 | | 3.20 | | 4.30 | | 5.43 | |
| Minimum | 4.50 | | 4.08 | | 0.87 | | 3.67 | | 3.49 | |
| Maximum | 29.82 | | 24.45 | | 25.67 | | 44.34 | | 44.43 | |
| **Females (N=59)** | | | | | | | | | | |
| Mean | | 9.67 | | 8.72 | | 2.47 | | 8.09 | | 8.65 |
| Standard deviation | | 4.66 | | 3.57 | | 2.91 | | 5.43 | | 7.08 |
| Minimum | | 4.55 | | 4.20 | | 0.88 | | 3.67 | | 3.60 |
| Maximum | | 29.82 | | 24.45 | | 20.80 | | 44.34 | | 44.43 |
| **Males (N=54)** | | | | | | | | | | |
| Mean | | 7.07 | | 6.87 | | 2.59 | | 6.91 | | 7.06 |
| Standard deviation | | 2.07 | | 1.97 | | 3.52 | | 2.47 | | 2.47 |
| Minimum | | 4.50 | | 4.08 | | 0.87 | | 3.89 | | 3.49 |
| Maximum | | 12.81 | | 14.23 | | 25.67 | | 13.20 | | 16.19 |

The average provisioning interval of the two days before capture differed significantly for females (t(58) = -3.096, p-value = 0.003) but not for males (t(53) = -1.618, p-value = 0.112). When we combine the t-test results with the average provisioning intervals, we see that the average provisioning interval of females before the capture had an increasing tendency, meaning a higher frequency of nest visits.
